# Free-running enzymatic oligonucleotide synthesis for data storage applications

**DOI:** 10.1101/355719

**Authors:** M.A. Jensen, P.B. Griffin, R. W. Davis

## Abstract

Here we present preliminary results for a method of oligodeoxynucleotide synthesis using terminal deoxynucleotidyl transferase (TdT) to generate mixed base homopolymer runs of defined sequence. In a process we have termed Free-Running Synthesis (FRS), we allow TdT to freely add multiple bases of a given monomer. With this method, homopolymeric runs can be used for writing DNA in a data storage capacity, where a stretch of A’s, for example, will be read as a single base, and the transitions between successive homopolymer runs encodes information. As a proof-of-concept, we demonstrate homopolymer additions of A, C and T onto the 3’ end of a 22 base initiator strand.

## Introduction

There is an exponential growth in data storage requirements, just as Moore’s Law is reaching its limits. Conventional storage paradigms write bit features onto planar media, and these are reaching fundamental limits. We are in need of an alternative means of storing data. Due to the compact size of the DNA molecule and its long term stability, it is a perfect storage unit, which will allow us to store and access exabytes of information in the same amount of space where currently only a few gigabytes can be stored.

Traditional solid-phase phosphoramidite chemistry for DNA synthesis suffers diminishing returns on strands > 100 bases (b). For example, synthesis of a 20 b at a coupling efficiency of 99.5%, will generate 90% full-length product (FLP); in contrast, for synthesis of 200 b at the same coupling efficiency, yield only 37% FLP (target yield = (coupling efficiency) ^n^, where n is the total number of bases per strand). Synthesis quality and yield for the phosphoramidite methodology are mostly limited by the hazardous chemicals used, such as trichloroacetic acid, pyridine, acetonitrile, THF, imidazole, tetrazole and dichloromethane (1).

Free Running Synthesis (FRS) using an enzyme, terminal deoxynucleotidyl transferase (TdT), is proposed here for generating sequence-defined, mixed base homopolymer runs. What makes synthesis using an enzyme particularly attractive for synthesizing oligonucleotides in an aqueous nature, is the length of the sequence that can be rapidly generated. Terminal transferase, specifically, is a template-independent polymerase, which is very efficient at adding 1000s of monomers to the 3’ end of ssDNA initiator (2).

Our objective is to provide a simple and effective method of FRS where TdT is allowed to bind an initiator strand, and build short homopolymer stretches of a given monomer (e.g., dATP, dCTP or dTTP) from the 3’ end. While TdT does not incorporate each base equally (e.g., A > C > T > G) (3), this is not a problem with FRS. By adjusting the concentrations and buffer components (Mg^2+^ / Co^2+^) for each dNTP, strand elongation will be controlled. It does not really matter that C is more efficient in polymerization than T; any stretch of dNTP (e.g., > 2 b) is interpreted as a single base for DNA data storage purposes with the transitions between short homopolymer regions encoding information. Also, with an efficient wash step following each reaction, excess monomers are flushed from the reaction to prevent contamination in downstream cycles.

It is also known that TdT is both a processive and distributive polymerase (1). Unlike template-dependent DNA polymerases that are namely processive in strand elongation, TdT will dissociate from the 3’ end of the initiator strand when the reaction environment is depleted of monomer substrates; TdT may then randomly bind to another initiator strand or persist free in solution until conditions are optimal for further strand elongation. By using a rapid washing step post-reaction, TdT is expected to remain attached to the initiator strand long enough for introduction of the next base in sequence. As the enzyme is maintained in a processive state, this will allow us to synthesize strands > 8000 b without having to replenish the enzyme or to wait for a chance interaction between TdT and the initiator strand.

## Materials and Methods

### Synthesis conditions

Reagents: 20 or 40 units TdT, 2.5 pm FRSi, 1000 pm dATP, dCTP or dTTP, 1 x buffer, 1 x CoCl_2_, water adjusted to 25 ul total reaction volume. Reaction conditions were as follows: 1) 20 or 40 units TdT were added prior (3 min) to dNTP to allow uniform distribution (4); 2) dNTP was then added and 3) reacted for 20 or 50 min; 4) reaction was stopped to prevent further dNTP incorporation (70 °C / 10 min); 5) sample was then passed through P-6 desalting column (Bio-Rad, 732-6200) to remove unincorporated dNTPs; 6) the eluent was lyophilized to 10 ul; and 7) replenished with fresh TdT, buffer, CoCl_2_ and next dNTP in the sequence. Steps 1-6 were repeated until the full sequence was synthesized.

Enzymatic synthesis of a specified strand length is determined by the monomer to initiator ratio (M/I) (2). For example, a 1000 b strand with 1 picomole (pm) initiator would require 1000 pm dNTP (homopolymer runs up to 8000 b have already been shown). And given that 1 unit of TdT has a nucleotide incorporation rate of 0.28 pm / sec (5), it would take < 1 hr. to synthesize 1000 b; this is equivalent to 3.6 sec / b compared to traditional phosphoramidite methodology for array synthesis at ~ 5 min / b (83 hr. to generate 1000 b). Also, current enzymatic synthesis strategies where TdT is pre-attached with unblocked dNTPs for sequence-defined stepwise addition, report cycle times between 20 and 30 sec (6).

### PCR, cloning and sequencing

For amplification of ACTC-TdT polymerized reaction product for Sanger sequencing: FRS reaction, 5 ul, Taq polymerase (NEB) 0.5 ul, Taq buffer (1x), dNTPs (NEB) final 0.2 mM, 5 pm forward primer (FRSi initiator), 5 pm reverse primer (T40), water added to 25 ul. Thermal-cycling conditions using an ABI Veriti: 95 °C/30 40* [95 °C/15s | 45 °C/30s | 68 °C/30s] 68 °C/3m. For sequence analysis, we poly-adenylated the 3’ end of each sample to introduce an anchor for T40 to anneal in PCR amplification; the initiator sequence (CTGAGTTCGCTACAGTGTACTC) was also used as a forward primer (Tm values, 50 °C and 55 °C, respectively). We followed the manufacturer’s recommended protocol (Invitrogen, TOPO TA Cloning Kit for Sequencing, 450030). Sequencing was performed by GenScript; for results in Fig. 1, PCR product was used; for results in Fig. 3, target samples were inserted into the pCR-4 TOPO vector, and plasmid purified.

**Figure 1.**
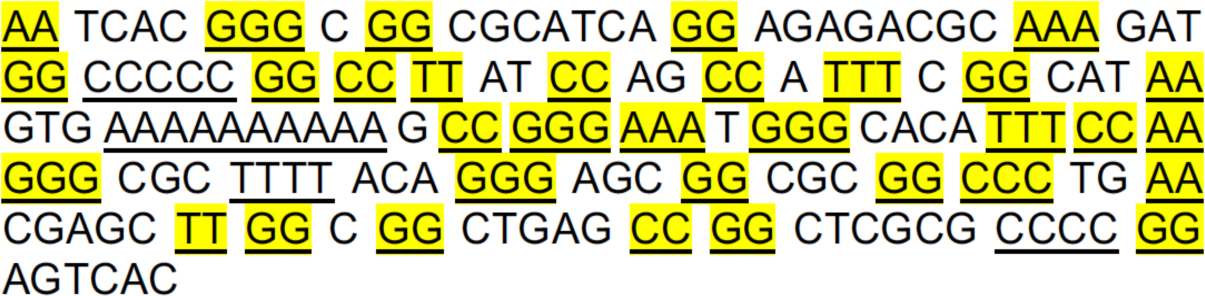
demonstrates an uncontrolled, free-running synthesis of single, doublet and triplet nucleotide incorporation (2 and 3 base additions highlighted in yellow). Synthesis conditions: initiator, 40 pm; 20 units of TdT (bovine, NEB), 2mM dNTP mix, 1× CoCl_2_, 1× TdT buffer, water added to a final volume of 20 ul, 37 °C / 60 min. Sample was PCR amplified and sequenced (see Material and Methods).

### Synthesis on super-paramagnetic beads (SPMB)

5′ Biotinylated FRS initiator strands (IDT) were coupled to streptavidin-coated MyOne SPMBs (Invitrogen) for 60 min in PBS at room temperature, 20 RPM. Samples were washed by magnetically collecting the SPMBs and removing the supernatant. This step was repeated three times with 200 ul PBS. For FRS with initiator bound to SPMB, reactions were placed in 1.5 ml tubes and incubated in a thermal mixing block at 37 °C and 95 °C (post-synthesis product cleavage from SPMB).

## Results and Discussion

For stepwise, enzymatic oligonucleotide synthesis, triphosphate nucleotide monomers would be generally blocked at the 3’ end to prevent homopolymeric tract formation. However, removal of the blocking moiety is the rate-limiting step of the cycle, which can add several minutes; this is necessary to assure complete separation of the blocking group, and that it is thoroughly washed to remove residual chemical deblocking agent and blocking group from the reaction (1). For the application of DNA data storage in particular, we have developed a method called Free-Running Synthesis (FRS), which allows us to build ssDNA without the requirement of using blocked nucleotides. TdT is most efficient when in a processive state where several nucleotide monomers are fed onto the strand, and TdT remains attached to the 3’ region. Based on the ratio of units TdT, monomer concentration (see Materials and Methods) and support loading of initiator strands, we can tailor approximately the number of homopolymeric dNTP additions that occur during each cycle (also contingent on buffer conditions, nucleotide (whether purine / pyrimidine)—TdT appears to have preference for incorporating some monomers over others, which affects their coupling efficiencies) (3). Presence of a divalent cation is also a critical determinant on the incorporation efficiency of a particular base (e.g., G > A > C > T (Mg^2+^), T > C > G > A (Co^2+^)); Zn^2+^ acts as an allosteric co-factor in addition to either Mg^2+^ and Co^2+^.

### Experiment 1: TdT polymerization with dNTP PCR mix

One of the first experiments we carried out was to see how TdT would behave randomly adding each base from an equimolar mix of dNTPs (Fig. 1).

Fig. 1 shows an evenly distributed heteropolymeric generation of ssDNA with nucleotide incorporation efficiency at G > C > A > T, where doublet and triplets were interspersed. Homopolymer stretches > 3 bases were also observed for A and C dNTPs (underlined). This simply demonstrates TdT will add each of the four bases onto an initiator strand, though short homopolymer stretches were tolerated. It is also noted that TdT has a preference for dATP self-polymerization.

### Experiment 2: Consecutive base additions (defined sequence)

In this next experiment, we synthesized A + C and C + A onto an initiator strand.

In Fig. 2, we demonstrate mixed base homo-polymerization of A added to C-polymerized initiator, and C added to A-polymerized initiator. To confirm C + A, we separately polymerized C for 90 min (duration of the total reaction in L3 where C was added for 60 min and then A for 30 min); if A was only polymerizing from residual initiator not consumed by the first C addition, then we should see an extra band in L3 that is level with the band shown in L2. It also appears each reaction was carried out to completion as the initiator strand (L1) is not visible in any of the other lanes (L2 – L5). What it also interesting, but not surprising, is that polymerization efficiency is different depending on whether A is added to C or C is added to A (L3 and L5) (7). For these reaction conditions, the order of efficiency is A (130 b) > A + C (100 b) > C + A (60 b) > C (40 b); for each sample, ladder minus 20 b (initiator) is equal to the length of strand polymerized (right image).

**Figure 2.**
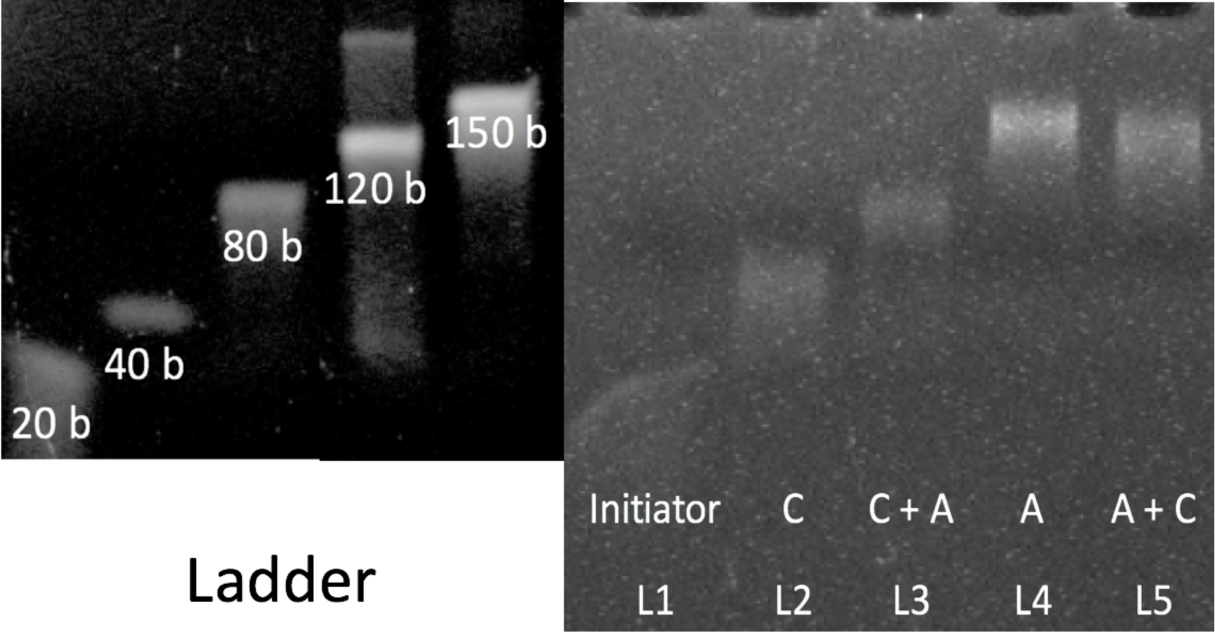
Comparison between addition of A onto C-polymerized initiator and addition of C onto A-polymerized initiator. Left image shows Ladder of single-stranded DNA, 20 b, 40 b, 80 b, 120 b and 150 b. Image on right, Lane 1 (L1): Initiator; L2: C added to initiator (60 min); L3: A added to C-polymerized initiator (30 and 60 min, respectively); L4: A-polymerized to initiator (90 min); L5: C added to A-polymerized initiator. All samples were stopped post-synthesis at 70 °C for 10 min; this time was included in the final incubation period since dNTP incorporation is expected to continue until TdT is fully inactivated as was observed with EDTA (4). Samples were run on 10% TBE PAGE (110 V / 60 min), and stained with EtBr. C and A were 1000 pm, initiator 2.5 pm; dATP = A; dCTP = C; based on preliminary rates of incorporation, we determined C added to initiator to near completion at 60 min, and A at 30 min (all initiator strand appeared to be consumed for each reaction).

### Experiment 3: Sequence analysis of a defined mixed base addition

Our final objective was to confirm the sequence identity of a defined multiple base addition onto an initiator strand. To minimize long, uncontrolled homopolymer runs that would cause self-annealing and secondary structure formation, we 1) adjusted reaction parameters to target synthesis of 10 – 50 bases (see Fig. 5), and 2) excluded G.

For this proof-of-concept, we have targeted synthesis of ACTC, which is initiated from the 3’ end of a general primer strand (for later PCR validation and sequencing analysis). Essentially, the steps were as follows: 1) TdT was mixed with buffer and initiator, and allowed to react at 37 °C for 3 min for pre-initiator-TdT binding, 2) the dNTPs were then added (e.g., dATP, 1000 pm) and then reacted for 20 min (or 50 min if using dCTP or dTTP), 3) the reaction was stopped by heat-inactivating the enzyme (70 °C for 10 min), 4) the reaction was desalted to remove unincorporated dNTPs, 5) the desalted product was lyophilized to 10 ul at 60 °C for 8 min, and then 6) steps 1 through 4 were repeated until the full-length strand was generated.

Fig. 3 demonstrates sequence-defined mixed base composition of homopolymer additions. This overlapping peak chromatogram is indicative of homopolymer runs during Sanger Sequencing. Furthermore, the short stretch of G’s after the poly A could be a sequencing artifact as there were no G’s introduced during FRS; this too, may be the result of polymerase slippage or mispriming from the complement strand.

**Figure 3.**
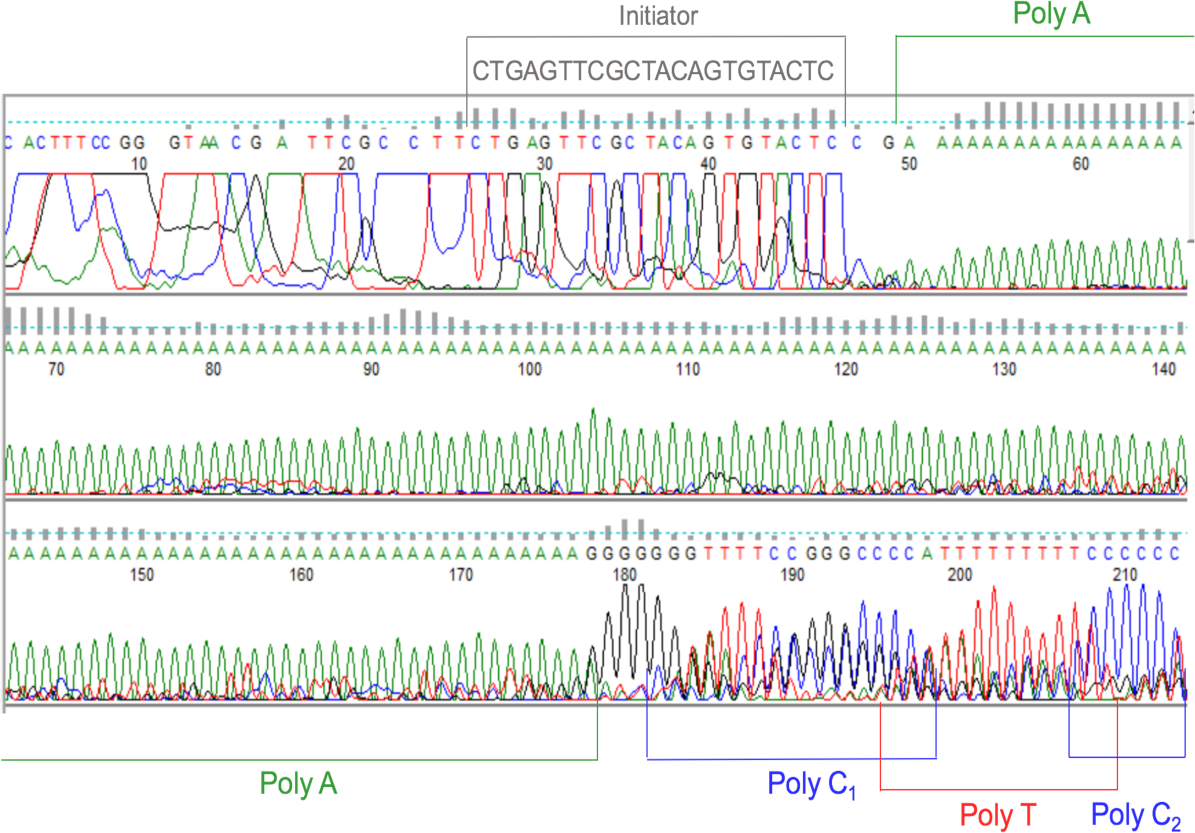
Sanger Sequencing trace chromatogram of a mixed base, homopolymer strand of defined sequence (CTGAGTTCGCTACAGTGTACTC—A_n_—C_n_—T_n_—C_n_, where n is any stretch of homopolymer bases). The initiator strand is loosely matched with base calls 26 through 47; poly A begins at base 50 through 177; poly C_1_, bases 182 through 198; poly T, 195 through 208; and poly C_2_, 207 through 213.

Also shown by the uninterrupted homopolymer stretches, the desalting step introduced between each cycle of ACTC FRS, was likely adequate enough to remove most of the unincorporated dNTPs. In a microfluidics channel, this would simulate the wash step.

## Considerations for optimal FRS

### Secondary structure formation and self-annealing during synthesis

For ease of use, we have omitted G from the sequence (Fig. 3) for the reason runs > 5 G nucleotides cause significant secondary structure formation; strands that knot at the 3’ end will preclude TdT binding and polymerization. However, to increase data storage alphabet, G can still be applied. For this, we suggest using either additives (e.g., betaine, DMSO, formamide, glycerol or trehalose) or substituting standard dGTP with 7-deaza dGTP (8).

Another drawback with producing homopolymeric, complementary base runs, is the increased likelihood of self-annealing. This will cause the strand to knot up and prevent TdT polymerization. For this, Pseudo-complementary triphosphate nucleotides could also be introduced (9).

### Reaction optimization

Because TdT does not incorporate the four standard bases equally, several synthesis parameters must be taken into consideration. For example, we noticed that it required 40 units TdT to polymerize approximately the same number of bases for dTTP as it did for dATP at only 20 units. dCTP also showed a similar increase in efficiency at 40 units. Buffer conditions would also need tailoring. For example, purines and pyrimidines each require a specific divalent cation to perform optimally (Mg^2+^ and Co^2+^, respectively); Zn^2+^ also acts as a cofactor to enhance polymerization efficiency for both cations.

Proofs-of-concept for this article were carried out under macro-scale synthesis conditions essentially for PCR and sequence validation. However, reaction times, units TdT, dNTP concentrations, and reaction volumes may be reduced orders of magnitude when applied to a microfluidics environment. For our application, we envisage a highly parallel synthesis platform for ssDNA generation at the attomole-scale for analysis on a nanopore sequencing device, and conversion from homopolymer base reads to digital output for DNA data storage with read-write capacity.

We also need to determine the average length of homopolymer stretches on a per cycle / reaction basis. By this we mean, that given a defined initiator density / substrate (e.g., pm of initiator / surface area of micro-well, bead or spot on array), we anticipate TdT will not be equally distributed over all initiator strands at any one time, even if the enzyme is in excess. Most enzymes will remain attached to the original initiator strand throughout synthesis; however, some will dissociate due to inadequate monomer concentration within proximity of the reaction site. As such, some strands will be longer than others. For example, we may target 10 b addition of dCTP, but may have 90% C10, 8% C9 and 2% C11. This is adequate information storage applications. If on the other hand we have 50% C10, 25% C2 and 25% C7, reaction parameters will require further optimization.

Fig. 4 compares how the ratio of TdT to initiator is critical to efficiently addressing each of the initiator strands in an equimolar amount (given an adequate monomer concentration). For example, if TdT < initiator, then TdT will only bind to a select number of initiator strands and continue to extend as long as there are monomers in the vicinity. Only when the concentration of monomers reaches a critical shortage, will the enzyme dissociate from the initiator and persist in solution until it contacts another initiator strand, and if the monomer concentration is sufficient for that reaction environment, will TdT bind the initiator and begin another round of polymerization.

**Figure 4.**
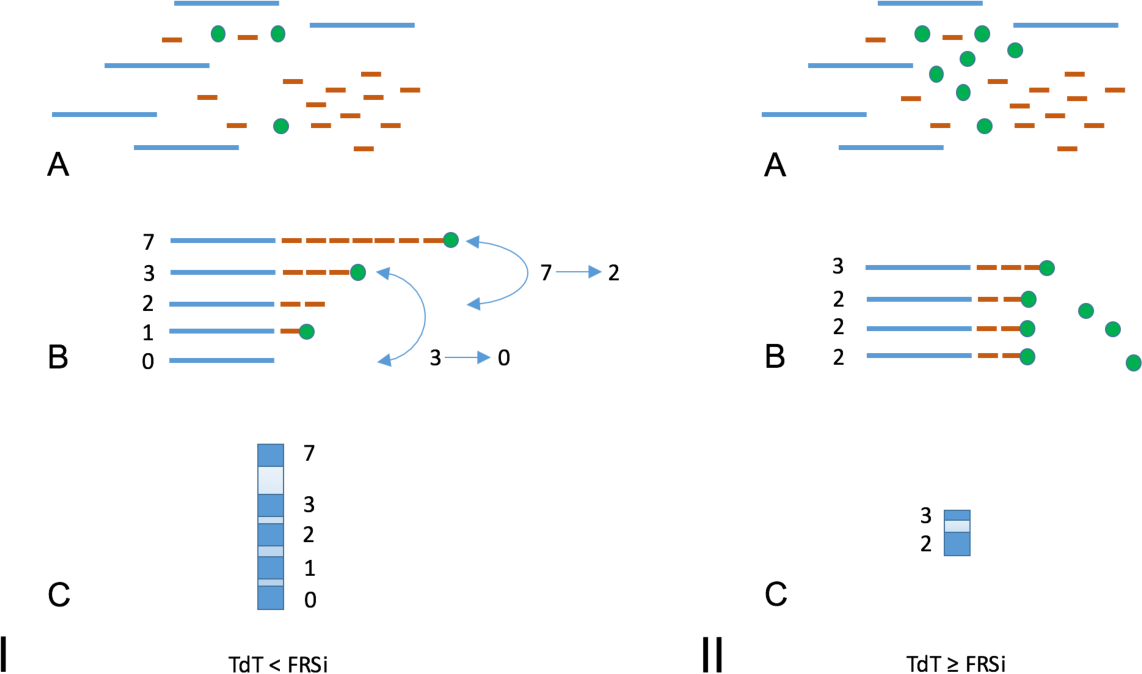
Theoretical effects of enzyme to initiator ratio on polydispersity. I: TdT < FRSi, II: TdT ≥ FRSi; A) relative reaction components; B) representation of polydispersity; C) representative final product band sizes. Blue bars, initiator; red dashes, dNTP; green dots, TdT; FRSi, Free-running synthesis initiator. Arrows in IB show TdT dissociation from parent strands, and attachment to daughter strands (2). FRSi, Free-Running Synthesis initiator.

As shown in Fig. 5, how much enzyme is available for the reaction has a significant baring on polymerization efficiency. For example, given the same initiator quantity (2.5 pm) and 1000 pm dATP monomer for 30 min, sample A2 with 40 units TdT shows a higher synthesis efficiency than sample A1 at only 20 units. The same is true for samples C2 and C3 (polymerization of dCTP) at 60 min. This corresponds with previous literature where Tang et al. found less polydispersity when the concentration of TdT was increased from 0.1 (0.05 unit of TdT / ul) to 2 (1 unit / ul), (4). Yield of initiator is also an important consideration in determining TdT polymerization efficiency. As shown in PAGE image left (Fig. 5), there is residual initiator remaining in sample A3 at 5 pm compared with A2 at 2.5 pm where it has been fully incorporated. If sample A3 was allowed to react with the next (different) base in sequence, left-over initiator would compete with the newly added dNTP, thus, adding to the sample’s polydispersity.

**Figure 5.**
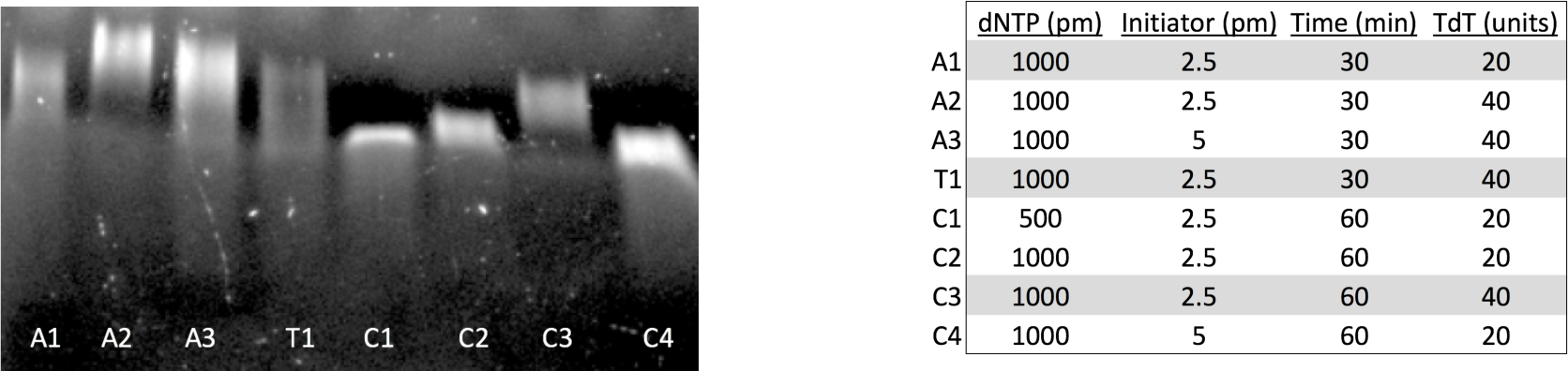
Effects of TdT concentration on polymerization efficiency. Image on left shows results for PAGE of various polymerization conditions detailed on the right. Conditions highlighted in grey were determined suitable for synthesis and sequencing ACTC, detailed in Experiment 3 above.

In Fig. 6, the concentration of monomer for a given reaction also dictates how well dNTPs will polymerize onto an initiator. It appears that a particular dNTP concentration is required for TdT to maximally access and polymerize monomers on the 3’ end of an initiator strand. For example, dTTP at 500 pm is not as efficient as synthesis with 1000 pm; however, increasing the reaction to 2000 pm had a negative effect on polymerization as shown by the difference in product migration (same with 5 and 10 pm initiator).

**Figure 6.**
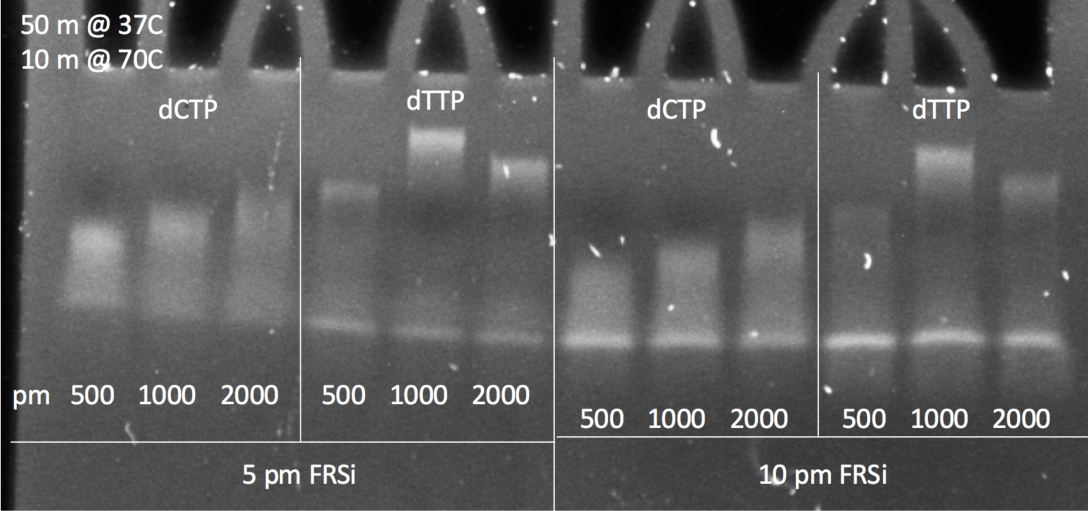
Effects of monomer concentration on polymerization efficiency. For each condition, a select monomer (either dCTP or dTTP) at 500, 1000 or 2000 pm, was polymerized on either 5 or 10 pm initiator strands for 50 min at 37 °C, followed by heat-inactivation for 10 min at 70 °C. Samples were run on 10% TBE PAGE at 90 V for 60 min.

## Conclusion

Here we show preliminary results for proof-of-concept of mixed base homopolymer ssDNA synthesis reactions (Fig. 2 and 3). Because all data collected for this phase of the project give a more general, relative product size comparison (10 – 100 b), the next phase will require us to focus in on DNA stretches with 2-10 b resolution; for this we will employ techniques such as autoradiography to measure single-base incorporation efficiencies at the sub-pm scale. While further validation may be necessary, we have so far demonstrated that we can 1) add dNTPs in a sequence-defined manner, 2) add base-specific monomers in homopolymer stretches per cycle, and 3) simulate an effective washing step between cycles by desalting each post-synthesis reaction to remove unincorporated dNTPs from further reacting in consecutive cycles. While these results are promising, certain limitations have precipitated that must be addressed to improve synthesis conditions. For example, some bases appear to have much higher incorporation efficiencies than others (e.g., A >> T or C) given similar reaction conditions. Though we lowered the incubation period for A to better normalize the rate of incorporation with C and T, this was still too unbalanced, and requires further optimization. Moreover, the Sanger method of sequencing is notorious for not accurately calling homopolymeric base runs, especially for A. To enable a scalable read strategy for information storage applications, we plan to introduce our samples on the nanopore device. Since the nanopore needs to simply distinguish between homopolymer transitions, the read requirements are much simpler than what is required to achieve single base resolution in biological applications. It is apparent too, that TdT maintains a preference for polymerizing certain bases; efficiency is also influenced by which base is already present on the 3’ end of the initiator strand. This is exemplified in Fig. 2 where A + C showed higher polymerization efficiency than C + A. To further tailor synthesis conditions, we have made several considerations regarding the ideal ratio of TdT to initiator to monomer. It is also apparent the units of enzyme required to fully address each of the initiator strands must be increased in order to prevent polydispersity. Also, too little monomer concentration will slow the reaction because TdT cannot access each dNTP fast enough to continue polymerization of an initiator strand; therefore, TdT will dissociate and persist in solution. By saturating the reaction with a select dNTP monomer, this will increase TdT catalysis; and when the reaction is stopped, whatever monomer remains in solution is washed clear to prevent contamination in downstream cycles.

Significant engineering optimization needs to be performed in order to quickly write short (2-mer or 3-mer) alternating homopolymer sequences useful for encoding information capable of exa-scale data storage applications. However, it seems that using unblocked natural nucleotides and the processive power of terminal transferase will provide a path to rapidly write sequences for large-scale information storage.

